## Supplemental Figures S1-S10 and Tables S1-S3 for "The solution structure of Dead End bound to AU-rich RNA reveals an unprecedented mode of tandem RRM-RNA recognition required for mRNA regulation"

### Figure S1

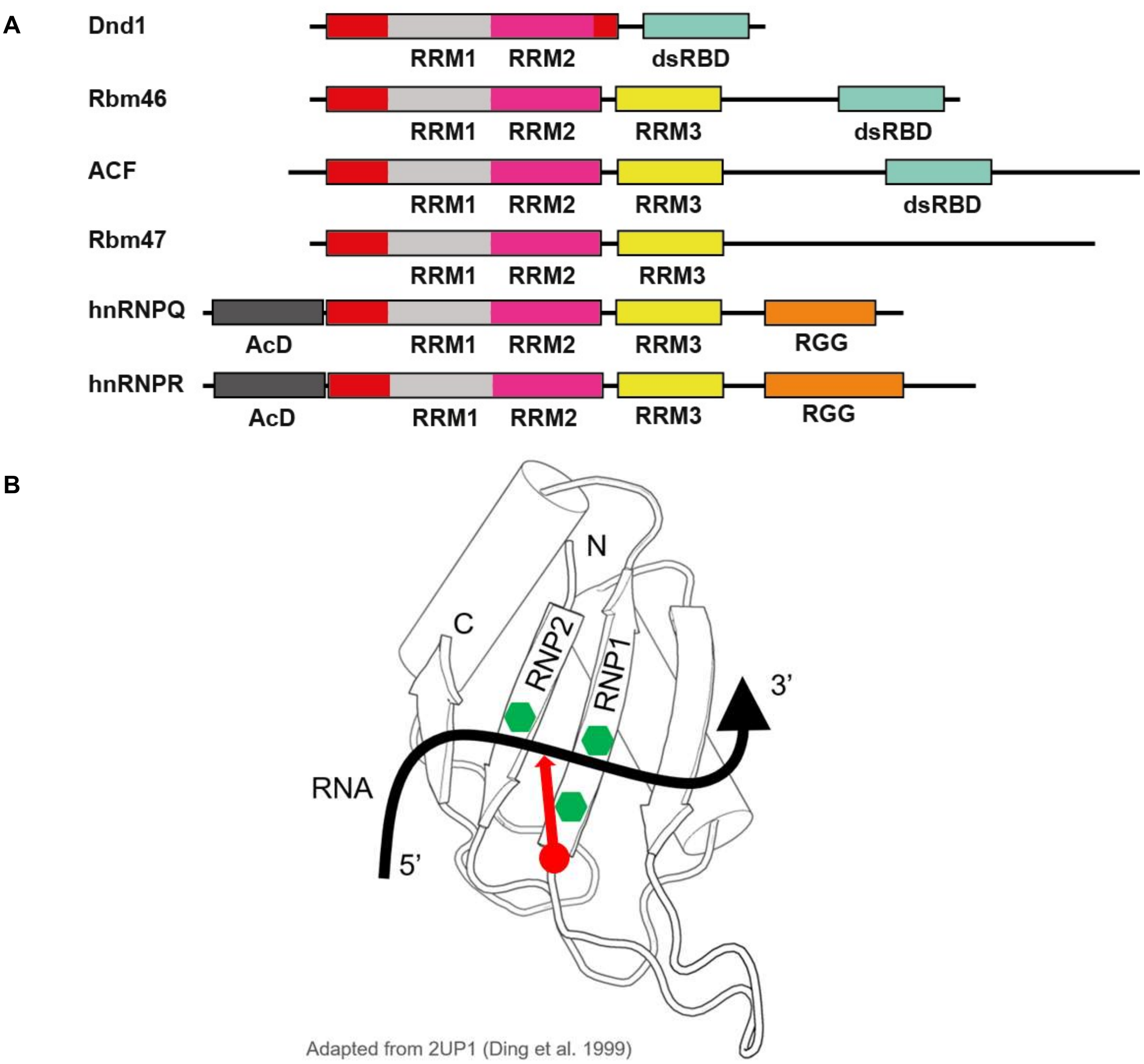


**Figure S1. Related to Figure 1. A) Domain structures of the hnRNPR-like family of proteins** that share sequence similarity to DND1 in its tandem RRMs, including the N-terminal extension. hnRNPQ and hnRNPR share a conserved N-terminal Acidic Domain (AcD) and C-terminal RGG repeats (Fig. S6). DND1, RBM46 and ACF share a dsRBD with low sequence conservation. Unique to DND1 is the lack of a third RRM, highly conserved in the other family members and lack of a canonical RNP RNA binding surface on RRM2. See Fig. S6 for sequence alignments. **B) Schematic view of a canonical RRM**. The canonical RRM fold is $\beta\alpha\beta\beta\alpha\beta$ folded into a four-stranded beta-sheet packed on top of two alpha-helices. The canonical RNA binding is of single-stranded RNA 5' to 3' over the beta-sheet starting at $\beta$4 towards $\beta$2. The central two beta-strands contain two conserved sequences RNP1 on $\beta$3 [R/K]-G-[F/Y]-[G/A]-[F/Y]-[I/L/V]-X-[F/Y] and RNP2 on $\beta$1 [I/L/V]-[F/Y]-[I/L/V]-X-N-L with three exposed aromatic sidechains (in green), two of which accommodate two bases by stacking interactions. In DND1 RRM1 these are F61 and Y102. A third aromatic sidechain (F100 in DND1 RRM1) usually stacks between the two ribose-moieties of these two nucleotides and an exposed long positively charged sidechain (in red) stabilizes the phosphate backbone between them. In DND1 RRM1 this is residue R98 that was mutated in the RIP assay (Figs. 1B, S3).

### Figure S2

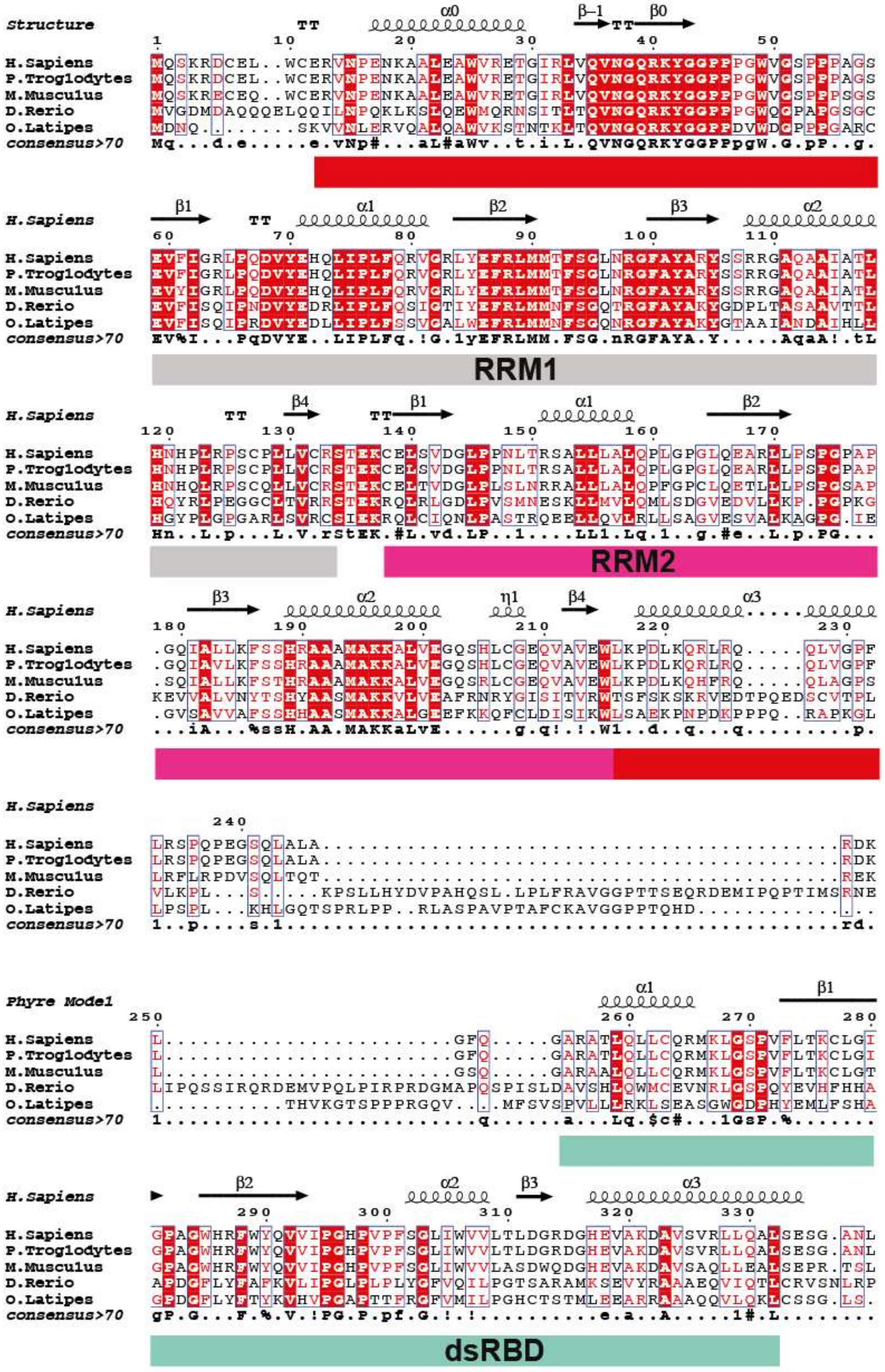
Structure
H.Sapiens
P.Troglodytes
M.Musculus
D.Rerio
O.Latipes
consensus>70
RRM1
RRM2
Phyre Model
dsRBD

**Figure S2. Related to Figure 1A. Sequence alignment of DND1** from human (Uniprot Q8IYX4, RefSeq NP919225.1), chimp (Uniprot H2QRM7, RefSeq XP_517978.2), mouse (Uniprot Q6VY05, RefSeq NP_775559.2), zebrafish (Uniprot Q7T1H5, RefSeq NP_997960.1) and medaka (Uniprot C7DQU6, RefSeq NP_001157988.1), prepared with the programs T-Coffee (Notredame et al. 2000) and ESPript 3.0 http://espript.ibcp.fr (Robert and Gouet 2014). Phylogenetic analysis reveals a high level of conservation between species within the RNA binding domains RRM1, RRM2 and dsRBD but also the N-terminal extension of RRM1 ($\alpha$0, $\beta$-1 & $\beta$0). Residues important for the interaction or discussed in the text are numbered according to the human protein sequence. Above the sequence alignment are the secondary structure elements based on either the tandem RRM12 structure presented in this paper or a model of the dsRBD calculated by the program PHYRE2 (Kelley et al. 2015) based on PDB entry 2l2n. Beta-turns are annotated by 'TT'. Below is a consensus sequence using the following criteria: **u**ppercase is identity, **l**owercase is consensus level > 0.7, **!** is any of I/V, **$** is any of L/M, **%** is any of F/Y, **#** is any of N/D/Q/E/B/Z. Colors and boxes are displayed on the alignment according to similarity scores calculated as described in (Robert and Gouet 2014): Red box, white character = Strict identity, Red character = Similarity higher than 0.7, Blue frame = Similarity of stretch of residues higher than 0.7.

**Figure S3**

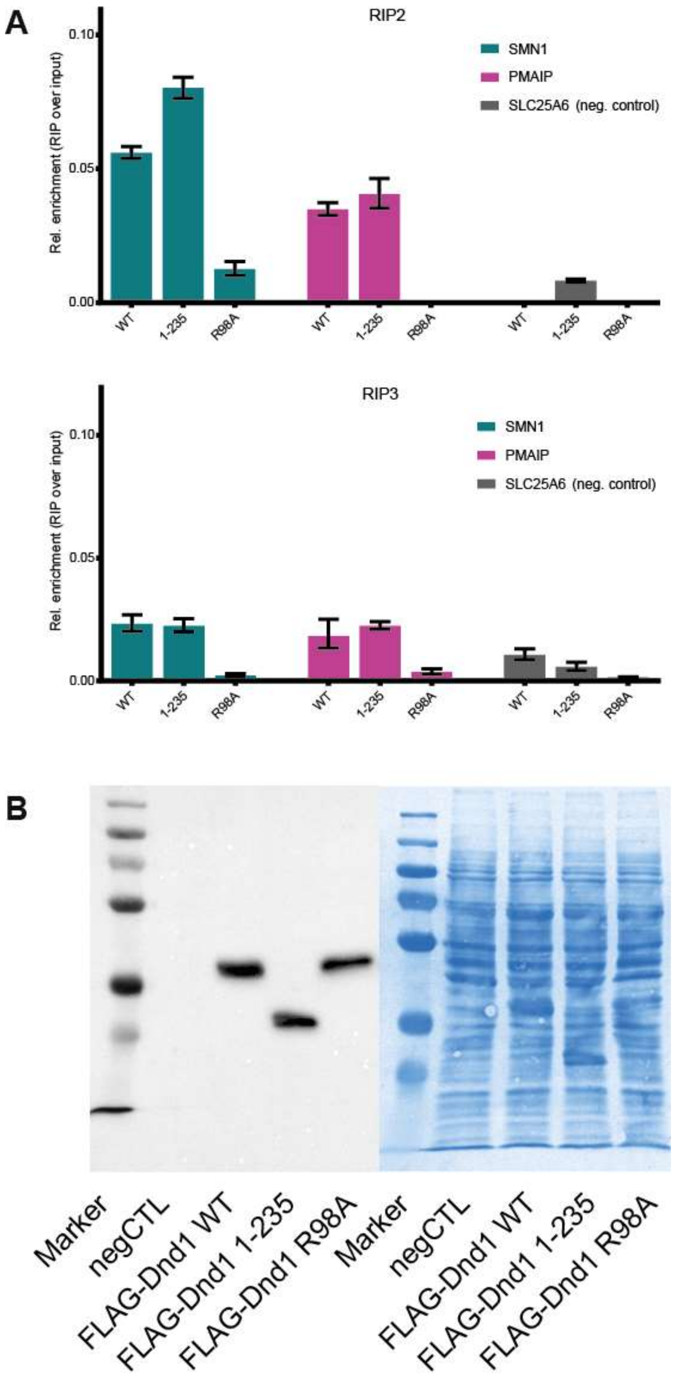


**Figure S3. Related to Figure 1B. A) RNA Immunoprecipitation** from HEK293T cells transiently expressing FLAG-tagged DND1 or its mutants followed by qPCR using primers for published DND1 targets and a negative control (Table S1). Representative data from two other independent experiment presented as relative enrichment over the input ($2^{-\Delta Ct}$). $\Delta$Ct is an average of (Ct [RIP] – (Ct [Input]) of technical triplicates with SD < 0.4. Error bars represent SD ($\Delta$Ct). **B) Expression in HEK293T cell culture.** Immunostaining with anti-FLAG antibody shows that DND1 and all mutants are well expressed in HEK293T cell culture. Western blot on the left, coomassie staining of the same gel on the right.

### Figure S4

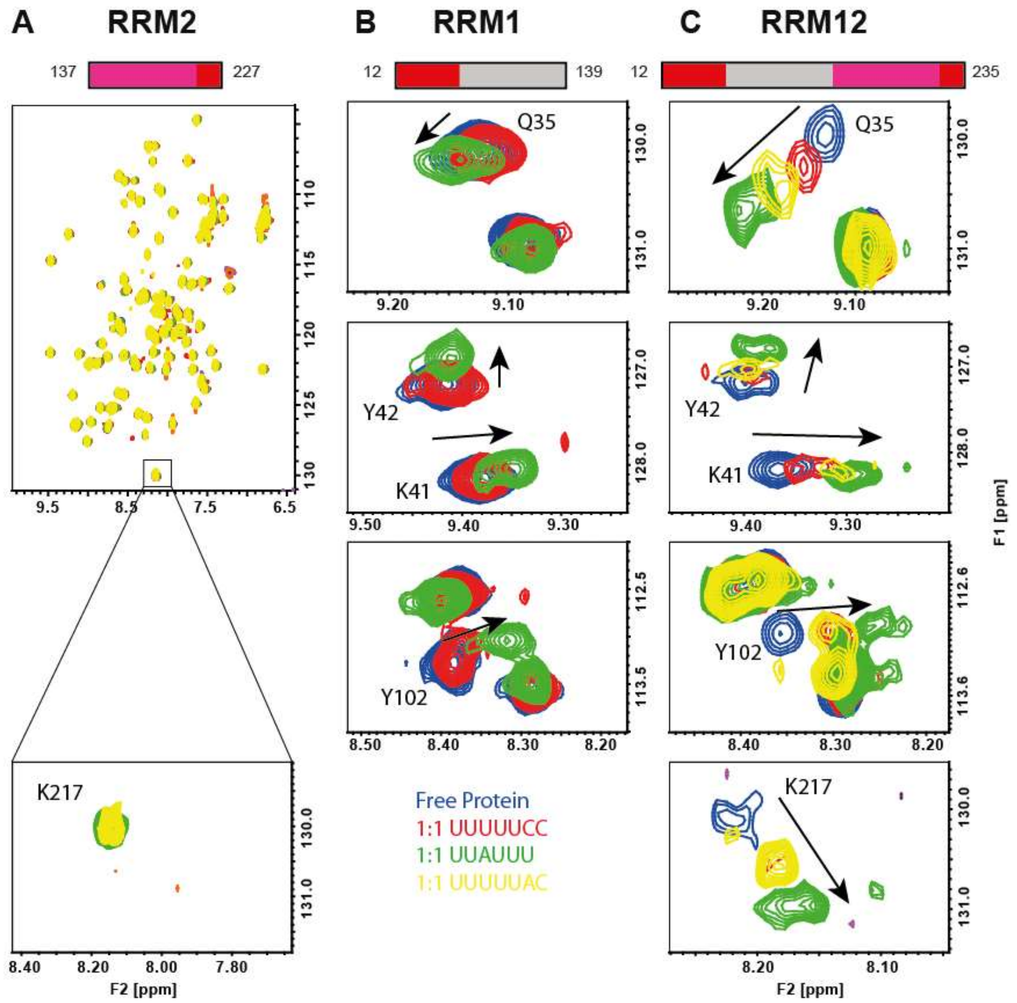


**Figure S4. Related to Figure 2. Finding the right protein construct and high affinity target for DND1-RNA structure determination using NMR titrations and ITC. A/B/C) Overlay of free and RNA-bound $^{1}H$-$^{15}N$-HSQC spectra of DND1 A) RRM2** includes a conserved C-terminal extension (137-227), full spectrum (top) and magnification of the individual cross-peak of the representative backbone amide of residue K217 in RRM2 alpha-helix 2 **B) RRM1** includes a conserved N-terminal extension (12-139), magnification of residues Q35, K41 and Y42 in the N-terminal extension and Y102 in RNP1. **C) RRM12** is the tandem RRM including both extensions (12-235), magnification of Q35, K41, Y42 and K217. The free protein spectra are shown in blue, spectra of a 1:1 molar equivalent complex with the oligos UUUUUCC, UUAUUU and UUUUUAC in red, green and yellow respectively. RRM2 alone does not bind any of the oligos. The arrows show increasing chemical shift perturbations upon introduction of an adenine into the oligo. Largest perturbations, indicative of higher affinity binding, are seen upon addition of an RNA oligo containing a central adenosine to the tandem RRM construct. Although the oligo UUAUUU including a central adenine can bind RRM1 alone, it shifts the signals originating from the bound residues to much lesser extent than when bound to the tandem RRM12, reflecting an increase in binding affinity upon addition of RRM2 as is confirmed by ITC in Figure 2 in the main text. A U-rich oligo not containing adenines at all (UUUUUCC) is not bound by RRM1 alone. This oligo, as well as an oligo with a peripheral adenine (UUUUUAC), is bound by the tandem RRM12 construct, although shifting the RNA bound residues to a lesser extent reflecting a weaker affinity. These titrations show that the affinity of DND1's tandem RRMs is highest to UUAUUU > UUUUUAC > UUUUUCC. NMR conditions: protein concentrations between 0.1 and 0.2mM, 298K, 750MHz. NMR buffer: 20mM MES pH 6.6, 100 mM NaCl, except for the RRM2 spectra recorded at 600MHz, in 25mM $K_2HPO_4$ / $KH_2PO_4$ buffer, 25mM NaCl, 2mM DTT pH6.3 at 303K.

**Figure S5**

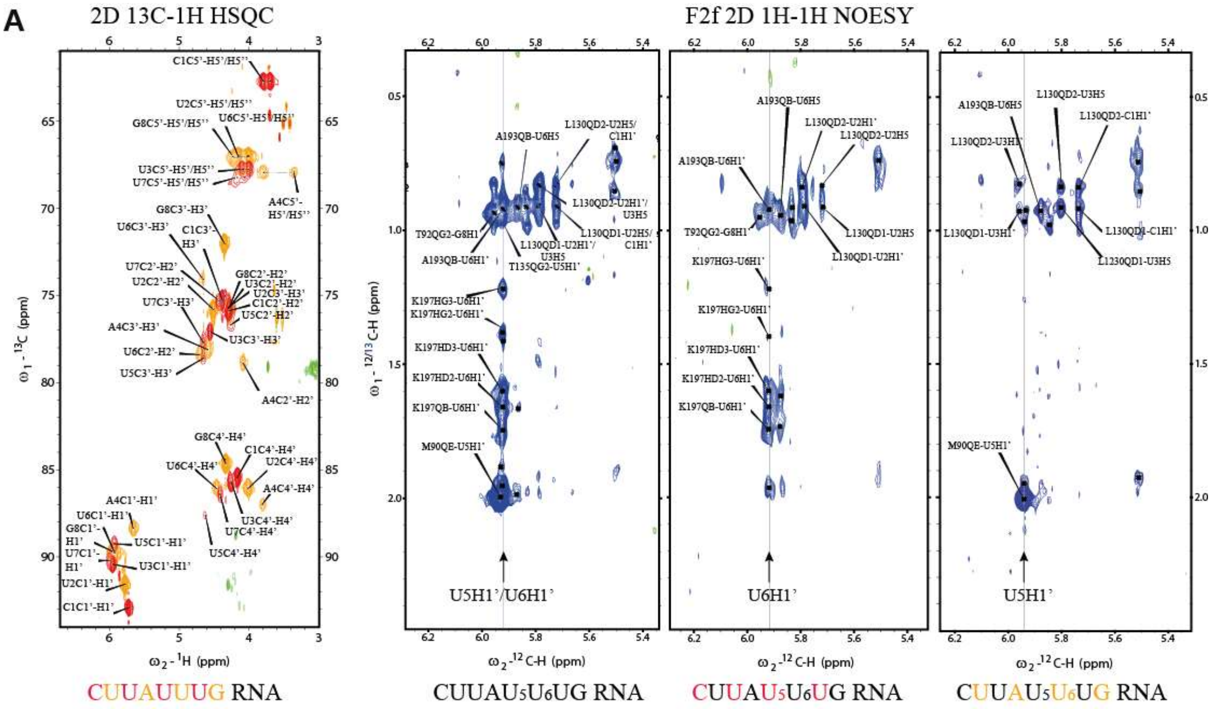
A
2D 13C-1H HSQC
F2f 2D 1H-1H NOESY
U5H1'/U6H1'
U6H1'
U5H1'
CUUAUUUG RNA
CUUAU5U6UG RNA
CUUAU5U6UG RNA
CUUAU5U6UG RNA

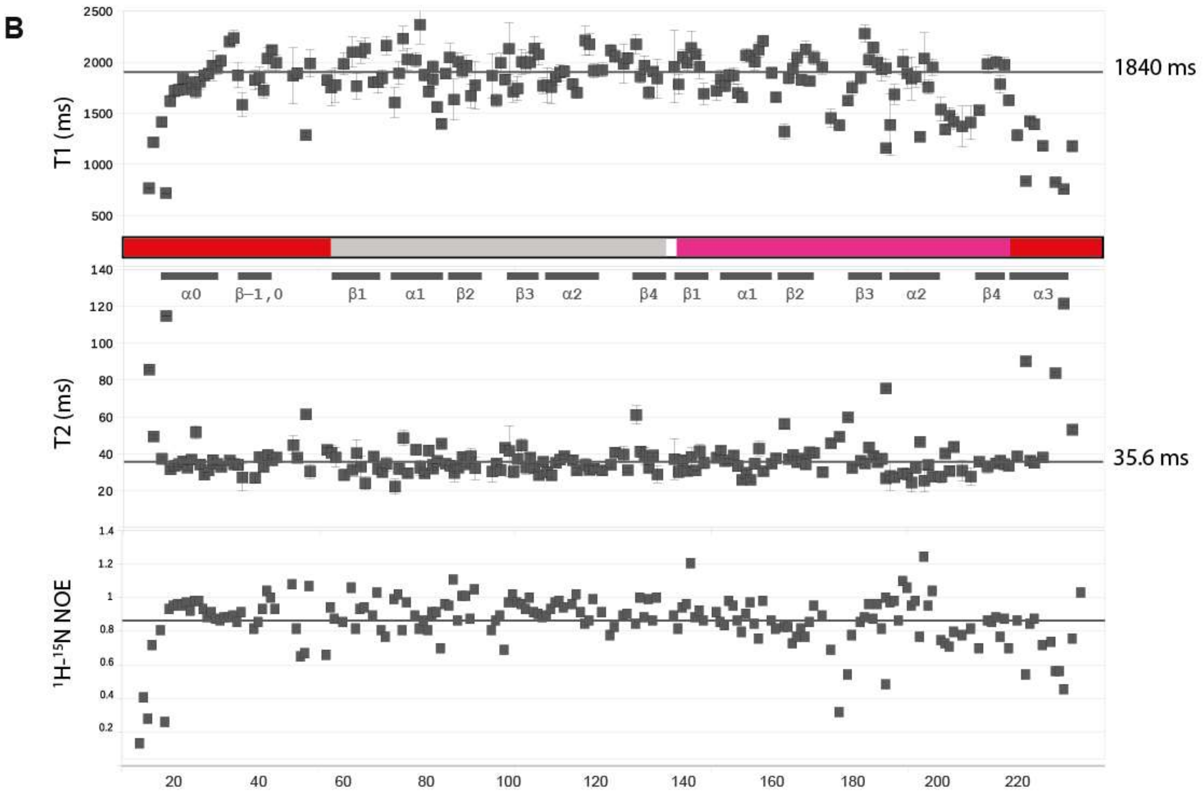
B
T1 (ms)
1840 ms
T2 (ms)
35.6 ms
1H-15N NOE
α0
β−1,0
β1
α1
β2
β3
α2
β4
β1
α1
β2
β3
α2
β4
α3

**Figure S5. Related to Figure 3. Selected NMR data DND1 RRM12-CUUAUUUG complex**

**A) Selective $^{13}$C-ribose labeling was essential for solving the structure of the DND1 tandem RRM – CUUAUUUG complex.** Solid phase synthesis using ribose-$^{13}$C labeled phosphoramidites facilitated the synthesis of short RNA oligos with alternating ribose moieties labeled with the NMR active nucleus $^{13}$C. Overlay of 2D $^{1}$H-$^{13}$C-HSQC spectra of CUUAUUUG RNA $^{13}$C ribose-labeled in either the 1$^{st}$, 3$^{rd}$, 5$^{th}$ and 7$^{th}$ (in red) or 2$^{nd}$, 4$^{th}$, 6$^{th}$ and 8$^{th}$ (in yellow) nucleotide position, in complex with the unlabeled tandem RRMs of DND1. Assignment of the heavily overlapped ribose protons is facilitated by resolution in the $^{13}$C dimension and reduction of assignment possibilities due to the selective labeling B) Excerpts of 2D NOESY spectra of complexes between the DND1 $^{13}$C$^{15}$N-labeled tandem RRMs and either unlabeled or selectively $^{13}$C-ribose labeled CUUAUUUG RNA, $^{13}$C filtered in the F2 dimension (F2f). Cross-peaks represent short distances (under 5 Ångstrom) between protons attached to unlabeled carbons only in the F2 dimension (in this case RNA base and unlabeled ribose) and protons attached to both $^{13}$C-labeled carbons (all protein and RNA labeled ribose) and unlabeled carbons (RNA base and unlabeled ribose) in F1. The arrow indicates the frequency at which both the $U_5$ and $U_6$ H1' protons resonate. By selectively labeling the $U_5$ ribose the cross-peaks to its H1' can be suppressed with a $^{13}$C-filter thus revealing NOEs specific to the $U_6$ ribose whereas a sample incorporating 13C-labeled $U_6$ ribose allows detection of NOEs to $U_5$, making unambiguous assignment of the short distances between the protein sidechains and the $U_5$ and $U_6$ H1' protons possible.

**B) Backbone dynamics of the DND1 tandem RRM-CUUAUUUG complex.** $^{15}$N-T1 and $^{15}$N-T2 relaxation rates and heteronuclear {$^{1}$H}-$^{15}$N NOEs (HetNOE) were measured by standard techniques described in the Method details and are plotted against the protein sequence where RRM1 is shown in grey, RRM2 in pink and the N- and C-terminal extensions are in red. Secondary structure elements are indicated below the sequence. Similar T1/T2 ratios for both RRMs including the N-terminal extension suggest isotropic motion, with the exception of the unfolded N- terminal tail, the C-terminal α3-helix and several loop residues. The average values of T2 and T1 of all residues with these exceptions indicated by solid lines are consistent with a globular protein of approximately 30 kDa. This is consistent with the size of a monomeric DND1 tandem RRM-CUUAUUUG complex determined by extrapolating from the correlation of $\tau$c and MW recorded at the same temperature (298K) and a similar field strength (Aramini et al., 2010). A rotational correlation time ($\tau$c) of 18ns was estimated using the formula

$$\tau_c \approx \left(\sqrt{\frac{6T_1}{T_2}-7}\right)/4\pi v_N$$

derived from Eq. 8 in Kay et al. (Kay et al. 1989) which considers only the contributions of J(0) and J(ωN) spectral density terms to T1 and T2 relaxation, neglecting higher frequency terms, and where $v_N$ is the $^{15}$N resonance frequency (in Hz).

### Figure S6

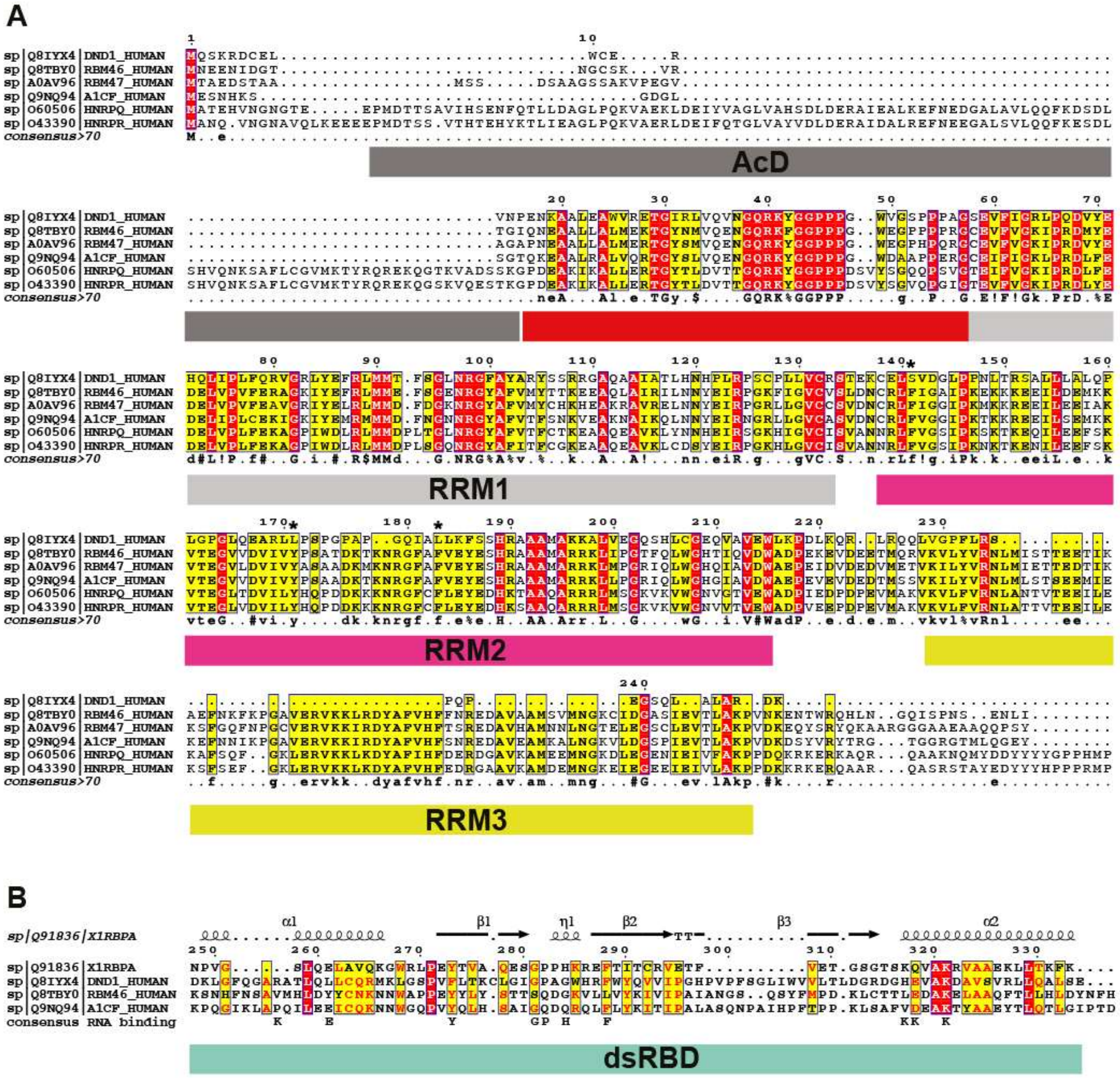

**Figure S6. Related to Figure S1. A) Sequence alignment of the N-terminal parts of the human hnRNPR-like family of RNA-binding proteins** DND1, RBM46, RBM47, ACF, hnRNPQ and hnRNPR. Uniprot identifiers are indicated in front of the protein names. Alignment was prepared as described in Fig. S2. Non-canonical RNP residues in DND1 are indicated with an asterix above the sequence. **B) Sequence alignment of the dsRBDs of the human hnRNPR-like family of RNA-binding proteins** DND1, RBM46 and ACF with the canonical second dsRBD of the protein XlRBPA with its secondary structure elements displayed above the sequence as found in PDBID 1DI2 (Ryter and Schultz 1998). Canonical dsRBD RNA binding residues are indicated below the sequence. Numbering according to DND1.

### Figure S7

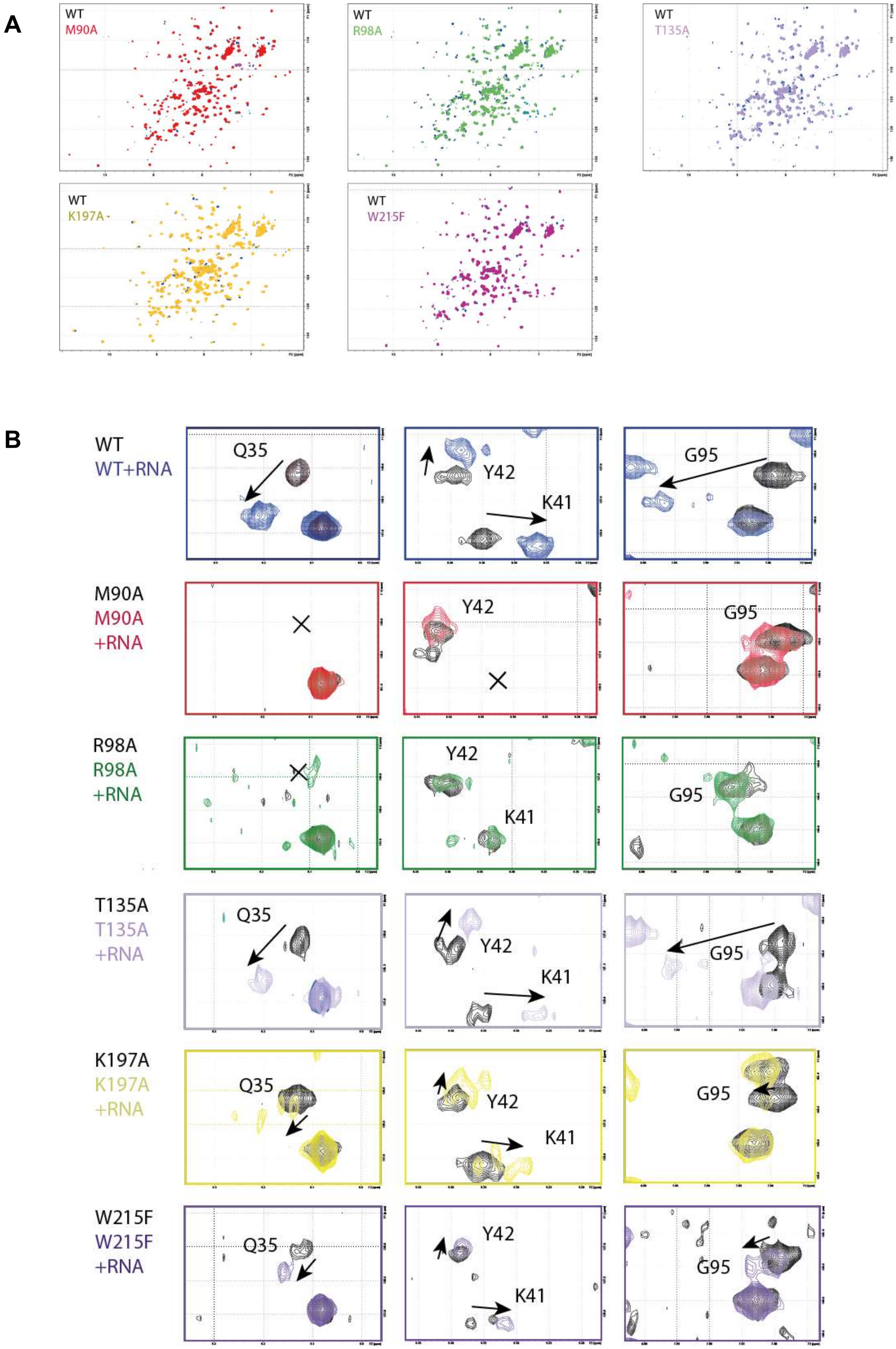
A
WT
M90A
WT
R98A
WT
T135A
WT
K197A
WT
W215F
B
WT
WT+RNA
Q35
Y42
K41
G95
M90A
M90A
+RNA
Y42
G95
R98A
R98A
+RNA
Y42
K41
G95
T135A
T135A
+RNA
Q35
Y42
K41
G95
K197A
K197A
+RNA
Q35
Y42
K41
G95
W215F
W215F
+RNA
Q35
Y42
K41
G95

**C**

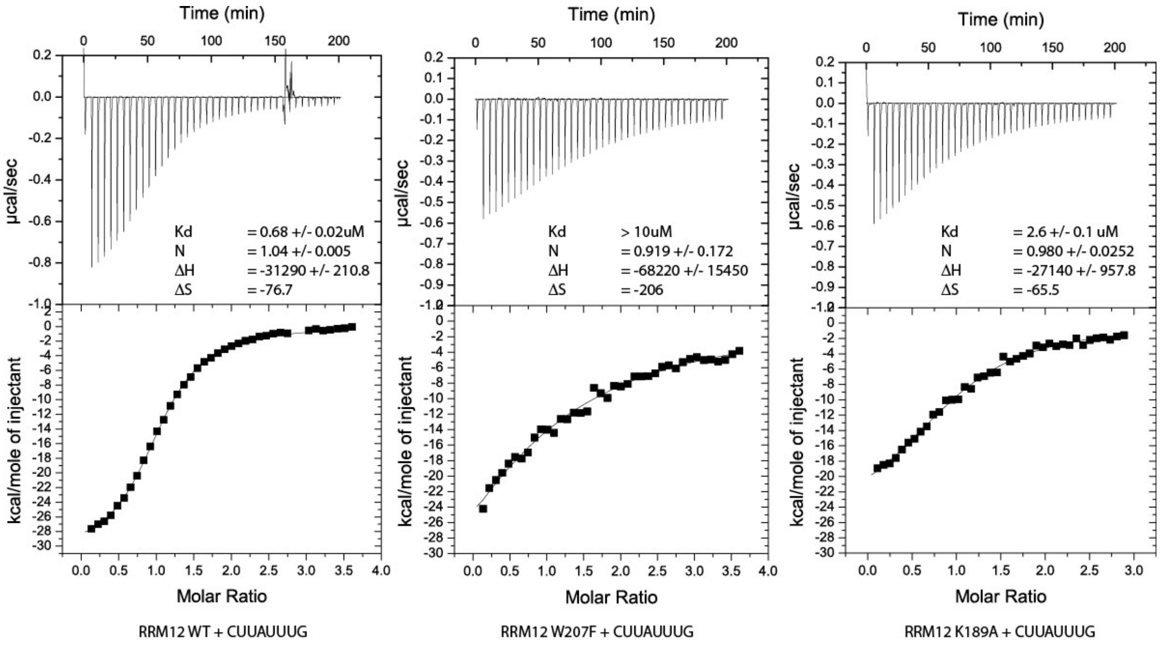


**Figure S7. Related to Figure 4. Mutational analysis of DND1 tandem RRM binding to CUUAUUUG RNA** A) $^{1}$H-$^{15}$N HSQC spectra of the WT tandem RRMs of DND1 are overlaid individually with the mutants M90A, R98A, T135A, K197A and W215F and show that all these mutants are correctly folded B) Disappearance or diminishing of chemical shift perturbations upon titration of CUUAUUUG RNA to several mutants compared to the WT tandem RRMs is observed for several reporter residues. C) **ITC measurement of DND1's tandem RRMs** domain titrated with CUUAUUUG RNA. WT: N = 1.04 +/- 0.005; $K_D$ = 0.68 +/- 0.02 uM; ΔH = -31.3 +/- 0.2 kcal/mol; -TΔS = 22.9 kcal/mol; W207F: N=0.919 +/- 0.172; $K_D$ = > 10uM; ΔH = -68 +/- 15 kcal/mol; -TΔS = 61.4 kcal/mol; K189A: N=0.98 +/- 0.03; $K_D$ = 2.6 +/- 0.1 uM; ΔH = -27.1 +/- 1.0 kcal/mol; -TΔS = 19.5 kcal/mol.

### Figure S8

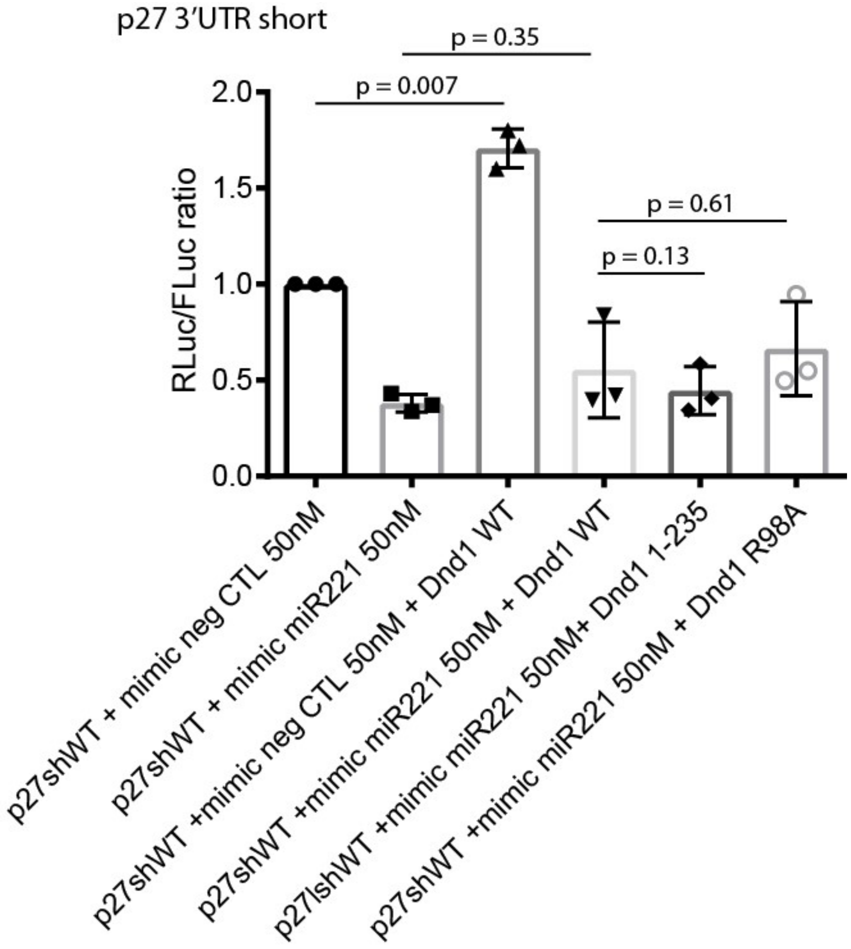


**Figure S8 Related to Figure 5. DND1 increases p27 3'UTR controlled reporter gene expression independent of the presence of a miR-mimic targeting this UTR or a negative control** A MiRIDIAN 221-3p miRNA mimic or negative control mimic was cotransfected into HEK293T with a short p27 UTR psiCHECK-2 luciferase construct and either WT or mutant FLAG-tagged DND1. Relative luciferase activity is the ratio between Renilla control and Firefly luciferases, adjusted to 1 for 100%. The results are represented as means and SD from three independent experiments. p-values obtained from two-tailed Welch's t-test.

### Figure S9

#### A) Proteins enriched in DND1 WT vs R98A RNA binding mutant interactome:

(DND1) ELAVL1, HNRNPL, PHAX, DHX30, PABPC1, SYNCRIP, MATR3, MOV10, MEX3A, TNRC6B, DHX9, HNRNPA2B1, LARP1, ILF3, SART3, HNRNPC, FUBP3

GO Biological Process 2018 ranked by EnrichR (Chen et al. 2013) combined score:

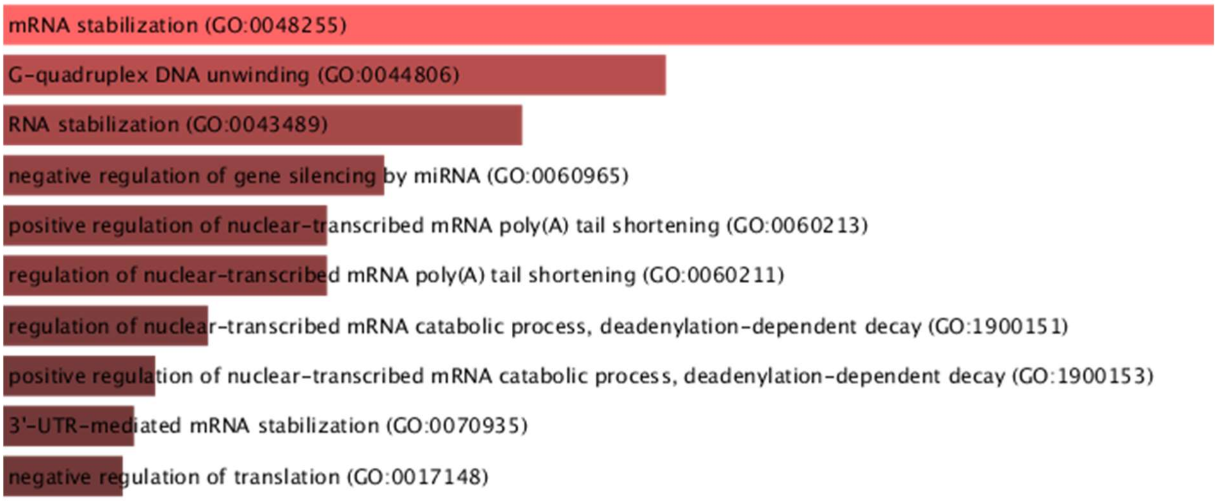


| Index | Name | P-value | Adjusted p-value | Odds Ratio | Combined score |
|---|---|---|---|---|---|
| 1 | mRNA stabilization (GO:0048255) | 2.282e-13 | 1.164e-9 | 202.02 | 5880.52 |
| 2 | G-quadruplex DNA unwinding (GO:0044806) | 0.00001602 | 0.006289 | 317.46 | 3505.25 |
| 3 | RNA stabilization (GO:0043489) | 4.129e-7 | 0.0002341 | 196.08 | 2882.36 |
| 4 | negative regulation of gene silencing by miRNA (GO:0060965) | 0.00003428 | 0.01093 | 222.22 | 2284.67 |
| 5 | positive regulation of nuclear-transcribed mRNA poly(A) tail shortening (GO:0060213) | 0.00004187 | 0.01257 | 202.02 | 2036.54 |
| 6 | regulation of nuclear-transcribed mRNA poly(A) tail shortening (GO:0060211) | 0.00004187 | 0.01187 | 202.02 | 2036.54 |
| 7 | regulation of nuclear-transcribed mRNA catabolic process, deadenylation-dependent decay (GO:1900151) | 0.00006917 | 0.01681 | 158.73 | 1520.46 |
| 8 | positive regulation of nuclear-transcribed mRNA catabolic process, deadenylation-dependent decay (GO:1900153) | 0.00009112 | 0.02114 | 138.89 | 1292.13 |
| 9 | 3'-UTR-mediated mRNA stabilization (GO:0070935) | 0.0001032 | 0.02290 | 130.72 | 1199.83 |
| 10 | negative regulation of translation (GO:0017148) | 1.204e-8 | 0.00001024 | 63.13 | 1151.22 |

GO Cellular Component 2018 ranked by EnrichR (Chen et al. 2013) combined score:

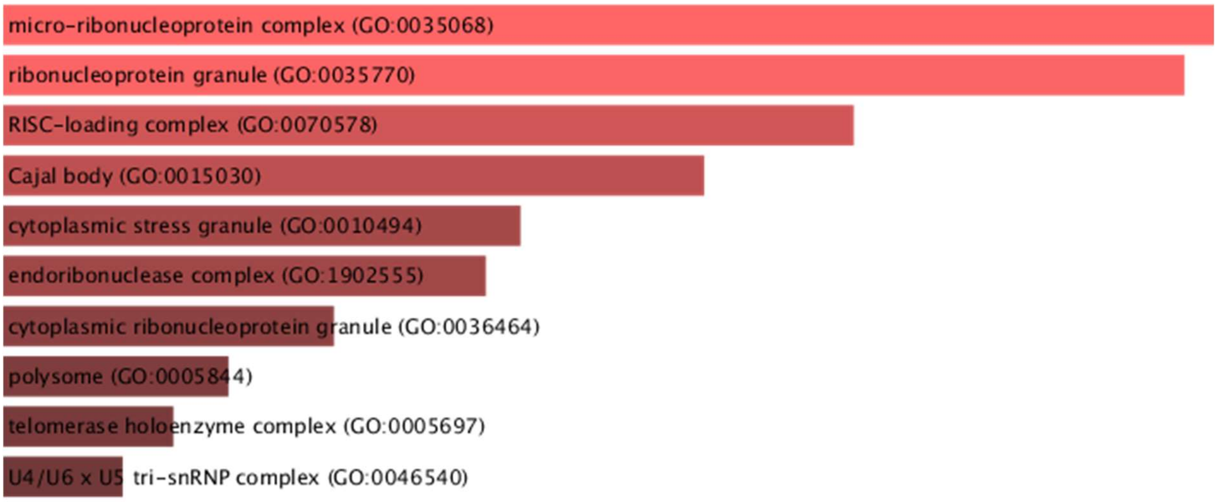


| Index | Name | P-value | Adjusted p-value | Odds Ratio | Combined score |
|---|---|---|---|---|---|
| 1 | micro-ribonucleoprotein complex (GO:0035068) | 0.006284 | 0.4004 | 158.73 | 804.73 |
| 2 | ribonucleoprotein granule (GO:0035770) | 6.958e-7 | 0.0003103 | 55.56 | 787.68 |
| 3 | RISC-loading complex (GO:0070578) | 0.008072 | 0.4500 | 123.46 | 594.98 |
| 4 | Cajal body (GO:0015030) | 0.0004219 | 0.06272 | 65.36 | 507.89 |
| 5 | cytoplasmic stress granule (GO:0010494) | 0.0006144 | 0.06851 | 54.20 | 400.81 |
| 6 | endoribonuclease complex (GO:1902555) | 0.01164 | 0.5768 | 85.47 | 380.62 |
| 7 | cytoplasmic ribonucleoprotein granule (GO:0036464) | 0.00001405 | 0.003133 | 26.14 | 292.11 |
| 8 | polysome (GO:0005844) | 0.001446 | 0.1290 | 35.27 | 230.64 |
| 9 | telomerase holoenzyme complex (GO:0005697) | 0.01962 | 0.7957 | 50.51 | 198.54 |
| 10 | U4/U6 x U5 tri-snRNP complex (GO:0046540) | 0.02227 | 0.7641 | 44.44 | 169.09 |

**B) Proteins enriched in DND1 WT vs 1-235 dsRBD truncation mutant interactome:**

(DND1) TRAP1, TNRC6B, HSPA1A, MATR3, MTHFD1, MEX3A, HSPA6, DDX17, HNRNPC

GO Molecular Function 2018 ranked by EnrichR (Chen et al. 2013) p-values

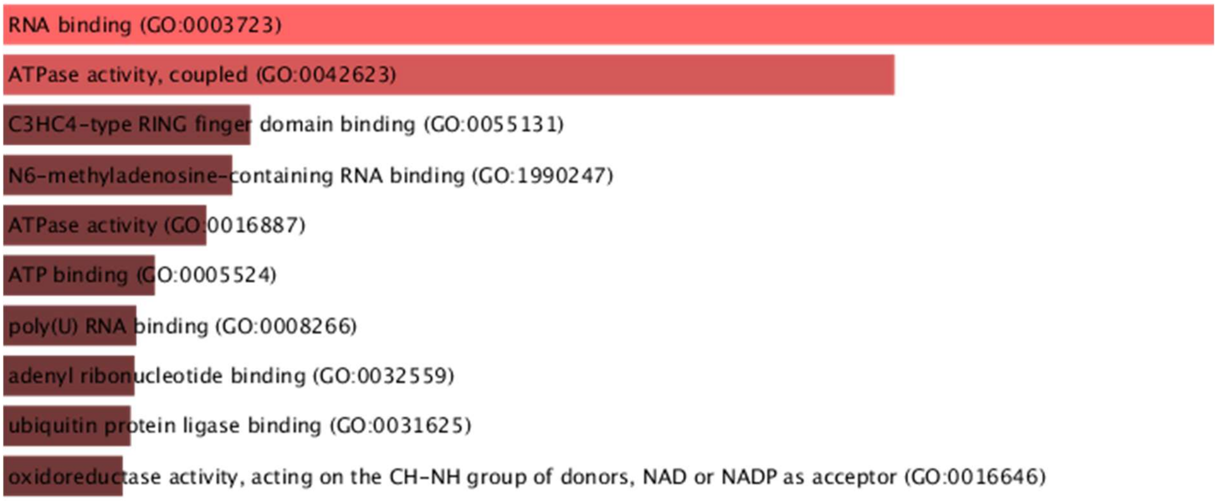


| Index | Name | P-value | Adjusted p-value | Odds Ratio | Combined score |
|---|---|---|---|---|---|
| 1 | C3HC4-type RING finger domain binding (GO:0055131) | 0.002997 | 1.000 | 333.33 | 1936.76 |
| 2 | N6-methyladenosine-containing RNA binding (GO:1990247) | 0.003495 | 1.000 | 285.71 | 1616.10 |
| 3 | ATPase activity, coupled (GO:0042623) | 0.00001178 | 0.006779 | 63.83 | 724.41 |
| 4 | poly(U) RNA binding (GO:0008266) | 0.007973 | 1.000 | 125.00 | 603.96 |
| 5 | oxidoreductase activity, acting on the CH-NH group of donors, NAD or NADP as acceptor (GO:0016646) | 0.008966 | 1.000 | 111.11 | 523.82 |
| 6 | poly-pyrimidine tract binding (GO:0008187) | 0.009957 | 0.9551 | 100.00 | 460.95 |
| 7 | AU-rich element binding (GO:0017091) | 0.01045 | 0.9255 | 95.24 | 434.37 |
| 8 | telomerase RNA binding (GO:0070034) | 0.01095 | 0.9001 | 90.91 | 410.42 |
| 9 | disordered domain specific binding (GO:0097718) | 0.01144 | 0.8781 | 86.96 | 388.73 |
| 10 | miRNA binding (GO:0035198) | 0.01194 | 0.8588 | 83.33 | 369.00 |

**Figure S9 Related to Figure 6. A) EnrichR (Chen et al. 2013) Gene ontology analysis of proteins enriched (t-test, FDR 10%) in WT vs mutant A: R98A; B: 1-235 DND1 HEK293T interactomes**

**Figure S10**

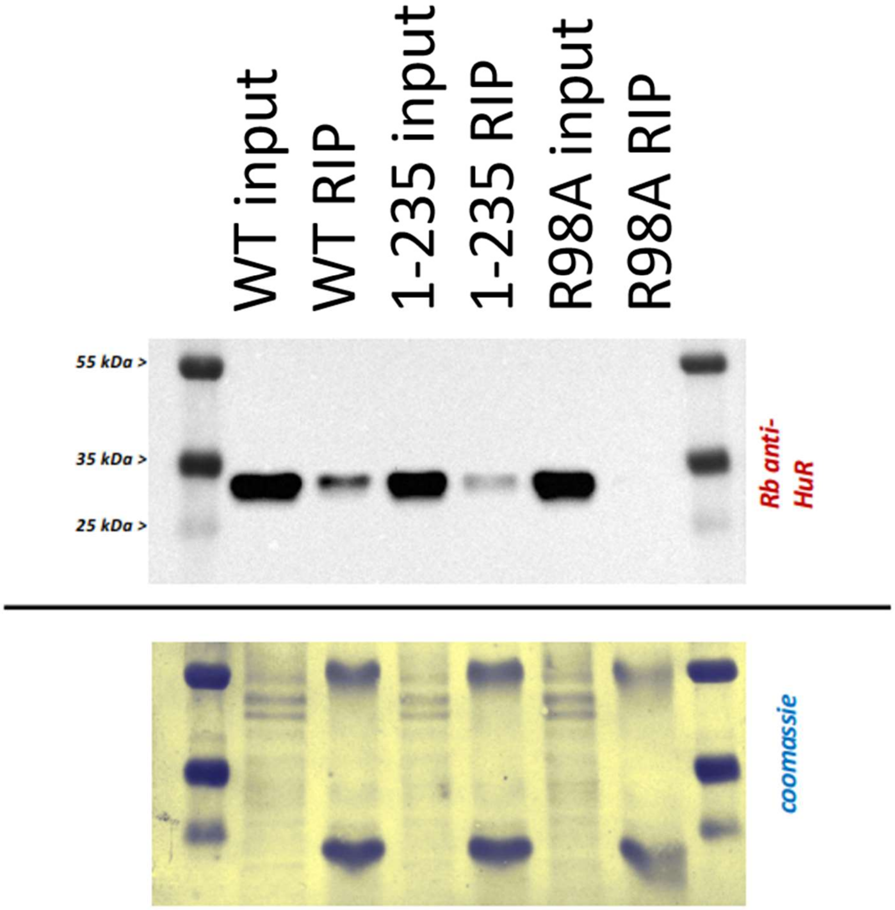


**Figure S10 Related to Figure 6. B) Western blot of ELAVL1/HuR from our WT, R98A and 1-235 DND1 transfected HEK293T pulldowns shows enrichment in WT and 1-235 dsRBD pulldowns only.**

**Table S1 related to Figures 1, S3 and Methods: Primers**

| Primer name | Primer sequence |
|---|---|
| Primers for qPCR | |
| SMN1_f | TGAGCTGTGAGAAGGGTGTT |
| SMN1_r | ACAGCCATGTCCACCAGTTAG |
| PMAIP_f | CTG GGC TAT ATA CAG TCC TCA AA |
| PMAIP_r | AGT GAC TAC AAC CTA CAT TAA GAC A |
| SLC25A6_f | CAG CCG ATT CCG TGT CTT GA |
| SLC25A6_r | GGG CTA GCG CTG AAG TAC AA |
| Primers for protein constructs | |
| hsDnd1_S1 | CGG GAT CCA TGC AGT CCA AGC GGG AT |
| hsDnd1_As353 | CGG GAA TTC CTA TCA TTA CTG TTT AAC CAT GGT ACC TGC CTC |
| hsDnd1_S12_BbsI_NcoI | CTA GAA GAC ACC ATG GAG AGG GTG AAT CCA GAG AAC AAG |
| hsDnd1_S136_BbsI_NcoI | CTA GAA GAC ACC ATG GAG AAG TGT GAG CTG AGC GTT GAC |
| hsDnd1_As139_Acc65I | GCT AGG TAC CTT ACT CAC ACT TCT CGG TGC TG |
| hsDnd1_As227_Acc65I | GCT AGG TAC CTT ACT GCT GGC GAA GTC GCT GCT TC |
| hsDnd1_As235_Acc65I | GCT AGG TAC CTT AGG ACC GCA AGA AGG GAC CCA C |
| Primers for protein mutations | |
| Mut_Dnd1_M90A_f | CGA GTT CCG CCT GGC GAT GAC CTT CAG |
| Mut_Dnd1_M90A_r | CTG AAG GTC ATC GCC AGG CGG AAC TCG |
| Mut_Dnd1_R98A_f | CAG CGG CCT GAA CGC CGG CTT CGC CTA TG |
| Mut_Dnd1_R98A_r | CAT AGG CGA AGC CGG CGT TCA GGC CGC TG |
| Mut_Dnd1_T135A_f | GTG TGC CGC AGC GCC GAG AAG TGT G |
| Mut_Dnd1_T135A_r | CAC ACT TCT CGG CGC TGC GGC ACA C |
| Mut_Dnd1_K197A_f | GCC ATG GCC AAA GCG GCC CTG GTG G |
| Mut_Dnd1_K197A_r | CCA CCA GGG CCG CTT TGG CCA TGG C |
| Mut_Dnd1_W215F_f | GTG GCT GTG GAG TTC CTC AAG CCA GAC C |
| Mut_Dnd1_W215F_r | GGT CTG GCT TGA GGA ACT CCA CAG CCA C |
| Primers for luciferase assay reporter constructs | |
| p27_sh3UTR_f_XhoI | GCT ACT CGA GGC CTC TAA AAG CGT TGG ATG |
| p27_sh3UTR_r_NotI | GCT AGC GGC CGC GTG CTA CAT AAA AGG TAA AAA CTA TAT ACA CAG |
| TERT_3UTR_f_XhoI | AAA ACT CGA GCG AGA GCA GAC ACC |
| TERT_3UTR_r_NotI | AAA AGC GGC CGC TTT TAC TCC CAC |
| TERT_mut_UAUUU_f | GCC CGG CTT TAT TTC CAC TCC CCA CAT AGG AAT AGT CCA TC |
| TERT_mut_UAUUU_r | GGG GAG TGG AAA TAA AGC CGG GCT CCT GGT GAG G |
| TERT_mut_UAUUU_ UUUU_UAUUU_f | GCC CGG CTT TAT TTT TTT TAT TTC CAC TCC CCA CAT AGG AAT AGT CCA TC |
| TERT_mut_UAUUU_ UUUU_UAUUU_r | GGG GAG TGG AAA TAA AAA AAA TAA AGC CGG GCT CCT GGT GAG G |

**Table S2. Related to Figures 2 and S4. Summary of *in vitro* protein-RNA interaction studies of the Dnd1 individual and tandem RRM domains with different RNA oligos.**

| oligo | RRM2 | RRM1 | | RRM12 | |
|---|---|---|---|---|---|
| | NMR | NMR | ITC (µM) | NMR | ITC (µM) |
| UUUUUCC | - | - | | + | > 100 |
| UUUUUAC | - | - | | ++ | > 35 |
| UUAUUU | - | + | | +++ | |
| CUUAUUUG | - | + | > 54 | +++ | 0.68 |

'-' means lack of binding in an NMR titration. Increasing numbers of '+' indicate binding with increasing affinity as indicated by increasing chemical shift perturbations of backbone amide proton NMR signals in a $^{1}H$-$^{15}N$ HSQC spectrum (see Fig. S4). NMR/ITC buffer: 20mM MES pH 6.6, 100 mM NaCl.

**Table S3. Related to Figures 3 and 4. Intra-RNA and Intermolecular restraints used to calculate the Dnd1 RRM12:CUUAUUUG complex ensemble**

**Intra-RNA restraints**

| Res1 | | Atom1 | Res2 | | Atom2 | Distance | Weight |
|---|---|---|---|---|---|---|---|
| 1 | RCYT | H6 | 1 | RCYT | H5 | 2.4 | |
| 1 | RCYT | H6 | 1 | RCYT | H1' | 4.5 | |
| 1 | RCYT | H6 | 1 | RCYT | H3' | 4.5 | |
| 1 | RCYT | H6 | 1 | RCYT | H2' | 3.5 | |
| 1 | RCYT | H6 | 1 | RCYT | H5' | 5 | |
| 1 | RCYT | H6 | 1 | RCYT | H5" | 5 | |
| 1 | RCYT | H6 | 1 | RCYT | H4' | 4.5 | |
| 1 | RCYT | H5 | 1 | RCYT | H5' | 5 | |
| 1 | RCYT | H5 | 1 | RCYT | H5" | 5 | |
| 1 | RCYT | H5 | 1 | RCYT | H3' | 5.5 | |
| 1 | RCYT | H5 | 1 | RCYT | H2' | 5 | |
| 1 | RCYT | H1' | 1 | RCYT | H5' | 5 | |
| 1 | RCYT | H1' | 1 | RCYT | H5" | 5 | |
| 1 | RCYT | H1' | 1 | RCYT | H4' | 4.5 | |
| 1 | RCYT | H1' | 1 | RCYT | H2' | 4.5 | |
| 1 | RCYT | H1' | 1 | RCYT | H3' | 4.5 | |
| 1 | RCYT | H2' | 1 | RCYT | H5' | 5 | |
| 1 | RCYT | H2' | 1 | RCYT | H5" | 5 | |
| 1 | RCYT | H2' | 1 | RCYT | H4' | 4.5 | |
| 1 | RCYT | H3' | 1 | RCYT | H5' | 4.5 | |
| 1 | RCYT | H3' | 1 | RCYT | H5" | 4.5 | |
| 1 | RCYT | H3' | 1 | RCYT | H4' | 4.5 | |
| 1 | RCYT | H3' | 1 | RCYT | H2' | 4.0 | |
| 1 | RCYT | H4' | 1 | RCYT | H5' | 4 | |
| 1 | RCYT | H4' | 1 | RCYT | H5" | 4 | |
| 1 | RCYT | H6 | 2 | URA | H5" | 6 | |
| 1 | RCYT | H1' | 2 | URA | H5' | 6 | |
| 1 | RCYT | H1' | 2 | URA | H5" | 6 | |
| 1 | RCYT | H3' | 2 | URA | H6 | 5 | |
| 1 | RCYT | H3' | 2 | URA | H5 | 5 | |
| 1 | RCYT | H3' | 2 | URA | H1' | 6 | |
| 1 | RCYT | H6 | 2 | URA | H6 | 5.0 | |
| 1 | RCYT | H5 | 2 | URA | H5 | 5.0 | |
| 1 | RCYT | H1' | 2 | URA | H6 | 5.0 | |
| 1 | RCYT | H4' | 2 | URA | H5" | 5 | |
| 2 | URA | H6 | 2 | URA | H1' | 4.5 | |
| 2 | URA | H6 | 2 | URA | H2' | 4.0 | |
| 2 | URA | H6 | 2 | URA | H3' | 4.5 | |
| 2 | URA | H6 | 2 | URA | H4' | 4.5 | |
| 2 | URA | H6 | 2 | URA | H5 | 2.4 | |
| 2 | URA | H6 | 2 | URA | H5' | 4.5 | |
| 2 | URA | H6 | 2 | URA | H5" | 4.5 | |
| 2 | URA | H5 | 2 | URA | H3' | 4.5 | |
| 2 | URA | H5 | 2 | URA | H2' | 4.5 | |
| 2 | URA | H1' | 2 | URA | H5' | 5 | |
| 2 | URA | H1' | 2 | URA | H5" | 5 | |
| 2 | URA | H1' | 2 | URA | H2' | 4.0 | |
| 2 | URA | H1' | 2 | URA | H4' | 4 | |
| 2 | URA | H1' | 2 | URA | H3' | 4 | |
| 2 | URA | H2' | 2 | URA | H3' | 3.5 | |
| 2 | URA | H3' | 2 | URA | H5' | 4.5 | |
| 2 | URA | H3' | 2 | URA | H5" | 4.5 | |
| 2 | URA | H3' | 2 | URA | H4' | 4.5 | |
| 2 | URA | H3' | 2 | URA | H5 | 4.5 | |
| 2 | URA | H4' | 2 | URA | H5' | 3.5 | |
| 2 | URA | H4' | 2 | URA | H5" | 3.5 | |
| 2 | URA | H4' | 2 | URA | H3' | 3.5 | |
| 2 | URA | H5' | 2 | URA | H5" | 3.5 | |
| 3 | URA | H6 | 3 | URA | H5' | 4.5 | |
| 3 | URA | H6 | 3 | URA | H5" | 5 | |
| 3 | URA | H6 | 3 | URA | H2' | 4 | |
| 3 | URA | H6 | 3 | URA | H3' | 4.5 | |
| 3 | URA | H6 | 3 | URA | H5 | 2.4 | |
| 3 | URA | H6 | 3 | URA | H1' | 4.5 | |
| 3 | URA | H1' | 3 | URA | H5" | 5 | |
| 3 | URA | H1' | 3 | URA | H5' | 5 | |
| 3 | URA | H1' | 3 | URA | H2' | 4.0 | |
| 3 | URA | H1' | 3 | URA | H4' | 4.5 | |
| 3 | URA | H1' | 3 | URA | H3' | 4.5 | |
| 3 | URA | H2' | 3 | URA | H5' | 4.5 | |
| 3 | URA | H2' | 3 | URA | H5" | 4.5 | |
| 3 | URA | H3' | 3 | URA | H5' | 4.5 | |
| 3 | URA | H3' | 3 | URA | H5" | 4.5 | |
| 3 | URA | H3' | 3 | URA | H2' | 4.0 | |
| 3 | URA | H3' | 3 | URA | H4' | 4.0 | |
| 3 | URA | H4' | 3 | URA | H5' | 3.5 | |
| 3 | URA | H4' | 3 | URA | H5" | 3.5 | |
| 3 | URA | H1' | 4 | RADE | H8 | 5 | |
| 3 | URA | H3' | 4 | RADE | H8 | 5 | |
| 3 | URA | H4' | 4 | RADE | H8 | 5 | |

| | | | | | | | |
|---|---|---|---|---|---|---|---|
| 3 | URA | H4' | 4 | RADE | H5' | 5 | |
| 3 | URA | H4' | 4 | RADE | H5" | 5 | |
| 4 | RADE | H2' | 3 | URA | H1' | 5 | |
| 4 | RADE | H8 | 4 | RADE | H1' | 4.5 | |
| 4 | RADE | H8 | 4 | RADE | H2' | 3.5 | |
| 4 | RADE | H8 | 4 | RADE | H3' | 4.5 | |
| 4 | RADE | H8 | 4 | RADE | H5' | 5 | |
| 4 | RADE | H8 | 4 | RADE | H5" | 5 | |
| 4 | RADE | H1' | 4 | RADE | H2' | 4.5 | |
| 4 | RADE | H1' | 4 | RADE | H4' | 4.5 | |
| 4 | RADE | H1' | 4 | RADE | H3' | 4.5 | |
| 4 | RADE | H3' | 4 | RADE | H5' | 4.5 | |
| 4 | RADE | H3' | 4 | RADE | H5" | 4.5 | |
| 4 | RADE | H3' | 4 | RADE | H4' | 4.5 | |
| 4 | RADE | H3' | 4 | RADE | H2' | 4.0 | |
| 4 | RADE | H1' | 5 | URA | H2' | 5.0 | |
| 4 | RADE | H1' | 5 | URA | H5 | 3.5 | |
| 4 | RADE | H1' | 5 | URA | H6 | 3.5 | |
| 4 | RADE | H3' | 5 | URA | H6 | 5 | |
| 4 | RADE | H8 | 5 | URA | H5 | 5 | |
| 5 | URA | H6 | 5 | URA | H1' | 4.5 | |
| 5 | URA | H6 | 5 | URA | H5 | 2.4 | |
| 5 | URA | H6 | 5 | URA | H2' | 3.5 | |
| 5 | URA | H6 | 5 | URA | H4' | 5 | |
| 5 | URA | H6 | 5 | URA | H3' | 4.5 | |
| 5 | URA | H5 | 5 | URA | H2' | 5 | |
| 5 | URA | H1' | 5 | URA | H2' | 4.0 | |
| 5 | URA | H1' | 5 | URA | H3' | 4.0 | |
| 5 | URA | H1' | 5 | URA | H5 | 5.5 | |
| 5 | URA | H2' | 6 | URA | H5 | 4.5 | |
| 5 | URA | H2' | 6 | URA | H6 | 4.5 | |
| 5 | URA | H3' | 6 | URA | H5 | 4.5 | |
| 6 | URA | H6 | 6 | URA | H1' | 4.5 | |
| 6 | URA | H6 | 6 | URA | H5 | 2.4 | |
| 6 | URA | H6 | 6 | URA | H2' | 4.5 | |
| 6 | URA | H6 | 6 | URA | H3' | 4.5 | |
| 6 | URA | H6 | 6 | URA | H4' | 6 | |
| 6 | URA | H6 | 6 | URA | H5' | 5 | |
| 6 | URA | H6 | 6 | URA | H5" | 6 | |
| 6 | URA | H5 | 6 | URA | H4' | 6 | |
| 6 | URA | H5 | 6 | URA | H2' | 5 | |
| 6 | URA | H1' | 6 | URA | H5' | 5 | |
| 6 | URA | H1' | 6 | URA | H5" | 5 | |
| 6 | URA | H1' | 6 | URA | H4' | 5 | |
| 6 | URA | H1' | 6 | URA | H3' | 5 | |
| 6 | URA | H1' | 6 | URA | H2' | 4.5 | |
| 6 | URA | H2' | 6 | URA | H5 | 4.5 | |
| 5 | URA | H3' | 6 | URA | H5' | 4.5 | |
| 5 | URA | H3' | 6 | URA | H5" | 4.5 | |
| 6 | URA | H1' | 7 | URA | H5" | 5.5 | |
| 7 | URA | H6 | 7 | URA | H2' | 4 | |
| 7 | URA | H6 | 6 | URA | H3' | 5 | |
| 7 | URA | H6 | 7 | URA | H5 | 2.4 | |
| 7 | URA | H6 | 7 | URA | H1' | 4.5 | |
| 7 | URA | H5 | 7 | URA | H2' | 5 | |
| 7 | URA | H5 | 6 | URA | H3' | 5.5 | |
| 7 | URA | H1' | 7 | URA | H2' | 4.5 | |
| 7 | URA | H1' | 7 | URA | H5 | 4.5 | 2 |
| 7 | URA | H2' | 7 | URA | H5' | 5 | |
| 7 | URA | H2' | 7 | URA | H5" | 4.5 | |
| 7 | URA | H2' | 8 | RGUA | H8 | 6 | |
| 7 | URA | H6 | 8 | RGUA | H1' | 4.5 | |
| 8 | RGUA | H8 | 8 | RGUA | H1' | 4.5 | |
| 8 | RGUA | H8 | 8 | RGUA | H3' | 4 | |
| 8 | RGUA | H8 | 8 | RGUA | H4' | 5 | |
| 8 | RGUA | H8 | 7 | URA | H4' | 6 | |
| 8 | RGUA | H1' | 8 | RGUA | H5" | 5 | |
| 8 | RGUA | H1' | 8 | RGUA | H5' | 5 | |

**Intermolecular restraints**

| Res1 | | Atom1 | Res2 | | Atom2 | Distance |
|---|---|---|---|---|---|---|
| 1 | RCYT | H1' | 130 | LEU | QD1 | 5 |
| 2 | URA | H6 | 130 | LEU | QD1 | 5 |
| 3 | URA | H6 | 130 | LEU | QD1 | 5 |
| 2 | URA | H6 | 130 | LEU | QD2 | 5 |
| 3 | URA | H6 | 130 | LEU | QD2 | 5 |
| 2 | URA | H5 | 130 | LEU | QD1 | 6 |
| 2 | URA | H5 | 130 | LEU | QD2 | 6 |
| 2 | URA | H1' | 130 | LEU | QD1 | 5 |
| 3 | URA | H5 | 130 | LEU | QD1 | 5 |
| 2 | URA | H1' | 130 | LEU | QD2 | 5 |
| 3 | URA | H5 | 130 | LEU | QD2 | 5 |
| 2 | URA | H2' | 130 | LEU | QD1 | 4 |
| 1 | RCYT | H2' | 130 | LEU | QD1 | 4 |
| 2 | URA | H2' | 130 | LEU | QD2 | 4 |
| 2 | URA | H2' | 130 | LEU | HG | 5 |
| 1 | RCYT | H2' | 130 | LEU | QD2 | 4 |
| 2 | URA | H3' | 130 | LEU | QD1 | 5 |

| | | | | | | |
|---|---|---|---|---|---|---|
| 2 | URA | H3' | 130 | LEU | QD2 | 6 |
| 3 | URA | H1' | 130 | LEU | QD1 | 5 |
| 3 | URA | H1' | 130 | LEU | QD2 | 5 |
| 3 | URA | H4' | 130 | LEU | QD2 | 5 |
| 3 | URA | H4' | 130 | LEU | QD1 | 5 |
| 3 | URA | H1' | 61 | PHE | HZ | 6 |
| 3 | URA | H5' | 64 | ARG | QG | 5 |
| 3 | URA | H5" | 64 | ARG | QG | 6 |
| 3 | URA | H5" | 64 | ARG | QD | 6 |
| 4 | RADE | H2 | 135 | THR | QG2 | 5 |
| 4 | RADE | H2 | 135 | THR | HB | 6 |
| 4 | RADE | H2 | 90 | MET | QE | 6 |
| 4 | RADE | H2 | 61 | PHE | QD | 6 |
| 4 | RADE | H2 | 102 | TYR | QE | 3.5 |
| 4 | RADE | H2 | 102 | TYR | QD | 4 |
| 4 | RADE | H8 | 61 | PHE | HZ | 6 |
| 4 | RADE | H8 | 61 | PHE | QE | 6 |
| 4 | RADE | H1' | 61 | PHE | QE | 6 |
| 4 | RADE | H1' | 61 | PHE | HZ | 6 |
| 4 | RADE | H1' | 100 | PHE | QD | 6 |
| 4 | RADE | H1' | 100 | PHE | QE | 6 |
| 4 | RADE | H2' | 100 | PHE | QE | 6 |
| 4 | RADE | H4' | 100 | PHE | QE | 5 |
| 4 | RADE | H5" | 100 | PHE | QE | 5 |
| 4 | RADE | H5" | 100 | PHE | QE | 5 |
| 4 | RADE | H1' | 90 | MET | QE | 5 |
| 4 | RADE | H3' | 100 | PHE | QE | 5 |
| 5 | URA | H6 | 137 | LYS | QD | 4 |
| 5 | URA | H5 | 137 | LYS | QD | 5 |
| 5 | URA | H1' | 137 | LYS | QD | 6 |
| 5 | URA | H6 | 137 | LYS | QG | 6 |
| 5 | URA | H5 | 137 | LYS | QG | 6 |
| 5 | URA | H5 | 135 | THR | QG2 | 6 |
| 5 | URA | H6 | 90 | MET | QE | 5 |
| 5 | URA | H1' | 102 | TYR | QR | 4.5 |
| 5 | URA | H4' | 102 | TYR | QR | 5 |
| 5 | URA | H1' | 100 | PHE | QE | 4.5 |
| 5 | URA | H1' | 189 | HIS | HE1 | 6 |
| 5 | URA | H4' | 100 | PHE | QE | 4.5 |
| 5 | URA | H2' | 100 | PHE | QE | 5.5 |
| 5 | URA | H4' | 90 | MET | QE | 5 |
| 5 | URA | H2' | 90 | MET | QE | 5 |
| 5 | URA | H1' | 90 | MET | QE | 3.5 |
| 6 | URA | H3' | 36 | VAL | QG1 | 6 |
| 6 | URA | H6 | 193 | ALA | QB | 5 |
| 6 | URA | H5 | 193 | ALA | QB | 4.5 |
| 6 | URA | H1' | 193 | ALA | QB | 3.5 |
| 6 | URA | H5 | 215 | TRP | HH2 | 4 |
| 6 | URA | H6 | 215 | TRP | HH2 | 6 |
| 6 | URA | H1' | 197 | LYS | HG3 | 5 |
| 6 | URA | H1' | 197 | LYS | HG2 | 5 |
| 6 | URA | H1' | 197 | LYS | HD3 | 4 |
| 6 | URA | H1' | 197 | LYS | HD2 | 4 |
| 6 | URA | H1' | 197 | LYS | QB | 4 |
| 6 | URA | H1' | 197 | LYS | QE | 3.5 |
| 6 | URA | H1' | 197 | LYS | HA | 6 |
| 6 | URA | H1' | 194 | MET | QE | 3.5 |
| 6 | URA | H4' | 194 | MET | QE | 4 |
| 6 | URA | H1' | 194 | MET | HA | 6 |
| 6 | URA | H2' | 197 | LYS | QE | 5 |
| 6 | URA | H4' | 197 | LYS | QE | 4 |
| 7 | URA | H4' | 36 | VAL | QG1 | 5 |
| 7 | URA | H3' | 37 | ASN | HD21 | 5 |
| 7 | URA | H5' | 36 | VAL | QG1 | 5 |
| 7 | URA | H5' | 39 | GLN | QG | 5 |
| 7 | URA | H5' | 36 | VAL | HB | 6 |
| 7 | URA | H5 | 92 | THR | QG2 | 5 |
| 7 | URA | H6 | 92 | THR | QG2 | 5 |
| 7 | URA | H4' | 197 | LYS | QE | 6 |
| 7 | URA | H5' | 197 | LYS | QE | 6 |
| 7 | URA | H5" | 197 | LYS | QE | 6 |
| 8 | RGUA | H8 | 93 | PHE | QE | 5 |
| 8 | RGUA | H8 | 92 | THR | QG2 | 6.0 |
| 8 | RGUA | H1' | 92 | THR | QG2 | 5.0 |
| 8 | RGUA | H3' | 92 | THR | QG2 | 6.0 |
| 8 | RGUA | H3' | 93 | PHE | QE | 5 |
